# Abundant non-A residues in the poly(A) tail orchestrate the mouse oocyte-to-embryo transition

**DOI:** 10.1101/2021.08.29.458077

**Authors:** Yusheng Liu, Hu Nie, Le-Yun Wang, Shuang Wu, Wei Li, Qi Zhou, Jiaqiang Wang, Falong Lu

## Abstract

Non-A (U, G, and C) residues can be added to the 5’-end, internal, and 3’-end positions of poly(A) tails of RNA transcripts^1–3^, and some of these have been shown to regulate mRNA stability^4, 5^. The mammalian oocyte-to-embryo transition (OET) relies on post-transcriptional regulation of maternal RNA, because transcription is silent during this process until the point of zygotic genome activation (ZGA)^6–9^. Although the regulation of mRNA translation by poly(A) tail length plays an important role in the mammalian OET, the dynamics and functions of non-A residues in poly(A) tails are completely unknown. In this study, we profiled the genome-wide presence, abundance, and roles of non-A residues during the OET in mice using PAIso-seq1 and PAIso-seq2^2, 10^, two complementary methods of poly(A) tail analysis. We found that non-A residues are highly dynamic in maternal mRNA, following a general pattern of beginning to increase at the MII stage, becoming highly abundant after fertilization with U residues in about half of poly(A) tails in 1-cell embryos, and declining in 2-cell embryos. We revealed that Btg4-mediated global maternal mRNA deadenylation created the substrates for U residue addition by Tut4/7 at their 3’-ends and further re-polyadenylation. In addition, G residues can be added by Tent4a/b. Finally, we demonstrate that G residues stabilize the modified mRNA, while the U residues mark maternal RNA for faster degradation in 2-cell mouse embryos. Taken together, these findings demonstrate that non-A residues are abundant and re-sculpt the maternal transcriptome to initiate zygotic development, which reveals the functional importance of the post-transcriptional regulation mediated by non-A residues in mRNA poly(A) tails.

## Introduction

The poly(A) tails found in most eukaryotic mRNAs have long been described as a chain of pure adenosine residues. However, recent studies have revealed that non-A (U, G, and C) residues can be added to the 3’ ends of the mRNA poly(A) tails^1–3^. In somatic cells, 3’-end U residues, catalyzed by TUT4/7, can promote rapid mRNA decay, whereas 3’-end G residues catalyzed by TENT4A/B can stabilize mRNA by protecting the poly(A) tails from rapid deadenylation^4, 5^. More recently, new poly(A) tail analysis methods based on the PacBio platform, including PAIso-seq (later called PAIso-seq1 to distinguish it from PAIso-seq2), PAIso-seq2, and FLAM-seq, reveal widespread non-A residues at the 5’-end and internal positions of the poly(A) tails in addition to the 3’-ends^2, 310^. These non-A residues in poly(A) tails (hereafter called non-A residues) are likely to constitute another layer of mRNA fate regulation.

A recent study indicated that poly(A) 3’-end U residues catalyzed by TUT4/7 are required for maternal mRNA decay following ZGA to ensure proper embryonic development in zebrafish^13^. Similar roles for global mRNA 3’-end U residues were also observed in *Xenopus* embryos^13^. The conditional knockout of *Tut4/7* in the germlines of both male and female mice impairs mRNA degradation through the 3’-end U residues, leading to failed spermatogenesis and oogenesis^11, 12^. Tut4/7 meditated 3’-end U residues are also required in maternal mRNA degradation after ZGA in mouse pre-implantation, as evidenced by the fact that siRNA mediated knockdown (KD) of *Tut4/7* in the zygote leads to maternal mRNA accumulation and impaired pre-implantation development^14^.

The oocyte-to-embryo transition (OET) in mammals is an essential process in development required for successful reproduction, and ranges from the meiotic resumption of oocytes at prophase of meiosis I to zygotic genome activation (ZGA) after fertilization^7–9^. Transcription is silent during the OET prior to ZGA^15^. These diverse biological processes are controlled by the maternal mRNA stored in oocytes, and are tightly regulated by post-transcriptional regulation mechanisms such as poly(A) tail length^7, 9, 16–19^. Deleting maternal *Btg4*, which encodes the adaptor of the CCR4-NOT poly(A) tail deadenylase complex, leads to failed OET with developmental arrest between the 1-cell to 2-cell stages due to defective deadenylation of maternal mRNA^20–22^. Similarly, deletion of *Cnot6l*, which encodes a catalytic subunit of the CCR4-NOT poly(A) tail deadenylase complex, also leads to failure of maternal mRNA clearance and embryo development^23^. In addition, increased mRNA tail length has been shown to positively regulate translation of several individual genes in mouse oocytes, including *Btg4*, *Cnot7*, *Tex19.1*, and *Oosp1*^20, 21, 24–27^. These findings indicate that poly(A) tail dynamics are essential to the functioning of the mammalian OET. However, the presence and role of non-A residues have not yet been studied during the mammalian OET.

Here, we applied two complementary methods, PAIso-seq1 and PAIso-seq2^2^, to comprehensively analyze the non-A residues during the OET in mice, including oocytes at the germ-vesicle (GV), metaphase I (MI), and metaphase II (MII) stages, as well as pre- implantation embryos at the 1-cell (1C), 2-cell (2C), and 4-cell (4C) stages. PAIso-seq1 has the advantage of sensitivity, with deep coverage for the low-input oocyte and embryo samples^2^, while PAIso-seq2 has the advantage of capturing all types of tails without any bias regarding polyadenylation or non-A residue status^10^. We found extensive non-A residues in maternal RNA, with variable abundance across stages. Analyzing the synthesis of these non-A residues provide a general picture that non-A residues are added to the deadenylated mRNA tails, which can then be further re-polyadenylated to generate non-A residues in 5’-end or internal parts of poly(A) tails. We revealed that the incorporation of non-A residues marks maternal mRNA, resulting in regulation of precise temporal stability, which is essential for the successful OET.

## Results

### Non-A residues in mRNA poly(A) tails are highly dynamic during the mouse OET

In order to explore global mRNA poly(A)-tail-mediated post-transcriptional regulation during the OET in mammals, we applied PAIso-seq1 and PAIso-seq2, two complementary methods for analyzing transcriptome-wide poly(A) tails^2, 10^, to analyze oocytes at the germ-vesicle (GV), metaphase I (MI), and metaphase II (MII) stages, as well as pre-implantation embryos at 1-cell (1C), 2-cell (2C), and 4-cell (4C) stages in mice (Fig. 1a). Two biological replicates demonstrated adequate reproducibility in gene expression and poly(A) tail length (Extended Data Fig. 1). Interestingly, we found that the number of poly(A) tails with non-A residues was highly dynamic during this process. The abundance increased at the MII stage, with a peak at the 1-cell stage, and decreased from the 2-cell stage to the 4-cell stage, where it reached a level comparable that of the GV stage (Fig. 1b, e). Among the U, C, and G residues, U residues were the most abundant, occurring in more than 40% of detected transcripts in mouse zygotes, followed by G residues, while C residues were the least abundant (Fig. 1b, e).

**Fig. 1.**
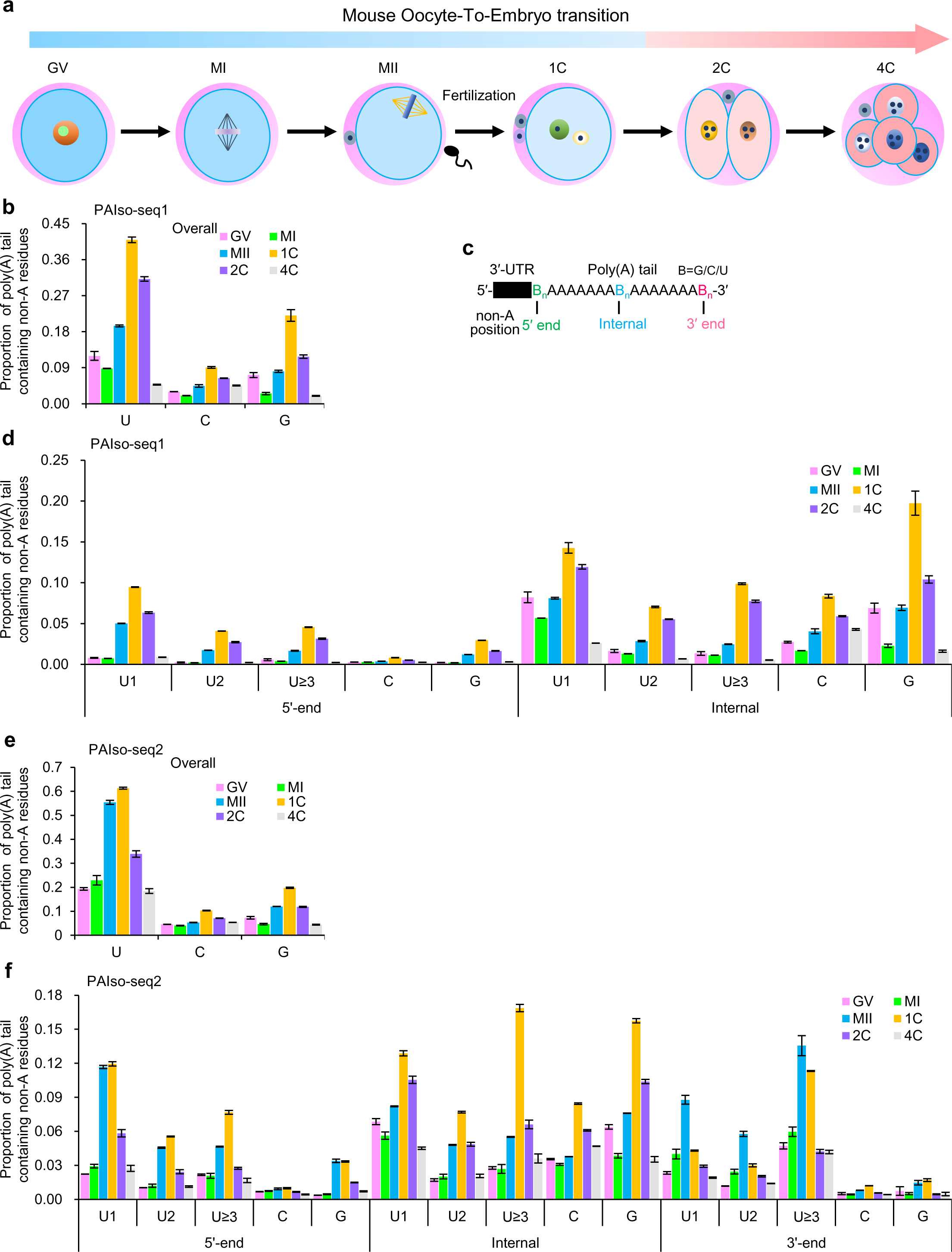
Non-A residues in mRNA poly(A) tails are highly dynamic during mouse OET. **a,** Schematic representation of the mouse oocyte-to-embryo-transition. GV, germ-vesicle oocyte; MI, metaphase I oocyte; MII, metaphase II oocyte; 1C, 1-cell embryo; 2C, 2-cell embryo; 4C, 4-cell embryo. **b, e,** Overall proportion of the transcripts containing non-A residues (U, C, or G) in samples at different stages measured with PAIso-seq1 (**b**) or PAIso-seq2 (**e**). Error bars represent the standard error of the mean (SEM) from two replicates. **c**, Diagram of the positions (5’-end, internal, and 3’-end) of non-A residues in poly(A) tails. **d, f,** Proportion of transcripts containing U, C, or G residues at the indicated positions (5’-end, internal, and 3’-end if available) of poly(A) tails in samples at different stages measured with PAIso-seq1 (**d**) or PAIso-seq2 (**f**). U residues were further divided according to the length of the longest consecutive number of Us (1, 2, and ≥3). Error bars represent the SEM from two replicates.

We then separated the non-A residues based on their relative positions in the poly(A) tails (Fig. 1c). U residues are often found in a consecutive manner. Therefore, we further separated U residues into three groups (U1, U2, U≥3) based on the maximum length of the consecutive U residues. The three groups of U residues as well as C and G residues showed overall similar dynamics in changes in abundance during mouse OET in both the 5’-end and internal parts of the poly(A) tails (Fig. 1d, f). The PAIso-seq1 and PAIso-seq2 datasets showed similar results, although PAIso-seq2 was additionally able to capture non-A residues at the 3’-ends.

No new transcription occurs during the OET before ZGA; therefore, the increased non-A residues can only be a result of processing of existing mRNA transcripts. For a given poly(A) tail containing 5’-end or internal non-A residues, the non-A residues must be added to the 3’-ends first, followed by further adenylation to produce a poly(A) tail with 5’-end or internal non-A residues. Interestingly, the pattern of U residues at the 3’-ends differed somewhat from that observed in the 5’-end and internal parts: it increased at the MI stage and peaked at the MII stage, and then was reduced at the 1-cell stage (Fig. 1d, f). This suggests that massive short or fully deadenylated mRNA 3’ tails appear during oocyte maturation^28^, which can be uridylated to produce a high level of 3’-end U residues. These 3’-end U residues can then be re-polyadenylated to produce 5’-end or internal U residues. 5’-end or internal U residues start to increase at the MII stage, indicating that re-polyadenylation has already begun at the MII stage and increases greatly after fertilization. C and G residues begin to increase at the MII stage and are further increased after fertilization regardless of position at the 5’-end, internal parts, or 3’-end (Fig. 1d, f). The abundances of C and G residues in the internal parts are much higher than those in the 5’-end or 3’-end, which differ from that of U residues (Fig. 1d, f). These suggest that different enzymes and mechanisms control the incorporation of G and C residues versus U residues. In addition, these differences also suggest different functions for U, C, and G residues.

The increase in 5’-end and internal non-A residues was beginning from the MI to MII stages, and further increased after fertilization. This indicates that global mRNA re-polyadenylation begins in MII oocytes and occurs on an even larger scale after fertilization. Interestingly, a similar phenomenon has also observed in rats, pigs, and humans^29, 30^.

Together, both the PAIso-seq1 and the PAIso-seq2 data revealed the tremendous changes in non-A residues between stages during the mouse OET, especially for the 5’-end and internal U residues, which represented around 40% of the mRNA transcripts in mouse zygotes.

### Characteristics of non-A residues in the mouse OET

To further analyze the highly abundant and dynamic non-A residues, we examined them at the individual gene level. The proportion of transcripts containing U, C, or G residues for a given gene increased at the MII stage, peaked at the 1-cell stage, and then decreased from the 2-cell stage to a low level at the 4-cell stage (Fig. 2a and Extended Data Fig. 2a). The major ZGA occurs at the 2-cell stage in mice, before which new transcription is minimal. Separating the maternal mRNA and the zygotic mRNA at the 2-cell stage clearly demonstrated that the maternal mRNA contained a higher level of non-A residues than did the zygotic mRNA (Fig. 2b-d and Extended Data Fig. 2b-d), revealing that the post-transcriptional regulation based on non-A residues was predominantly occurring on maternal mRNA.

**Fig. 2.**
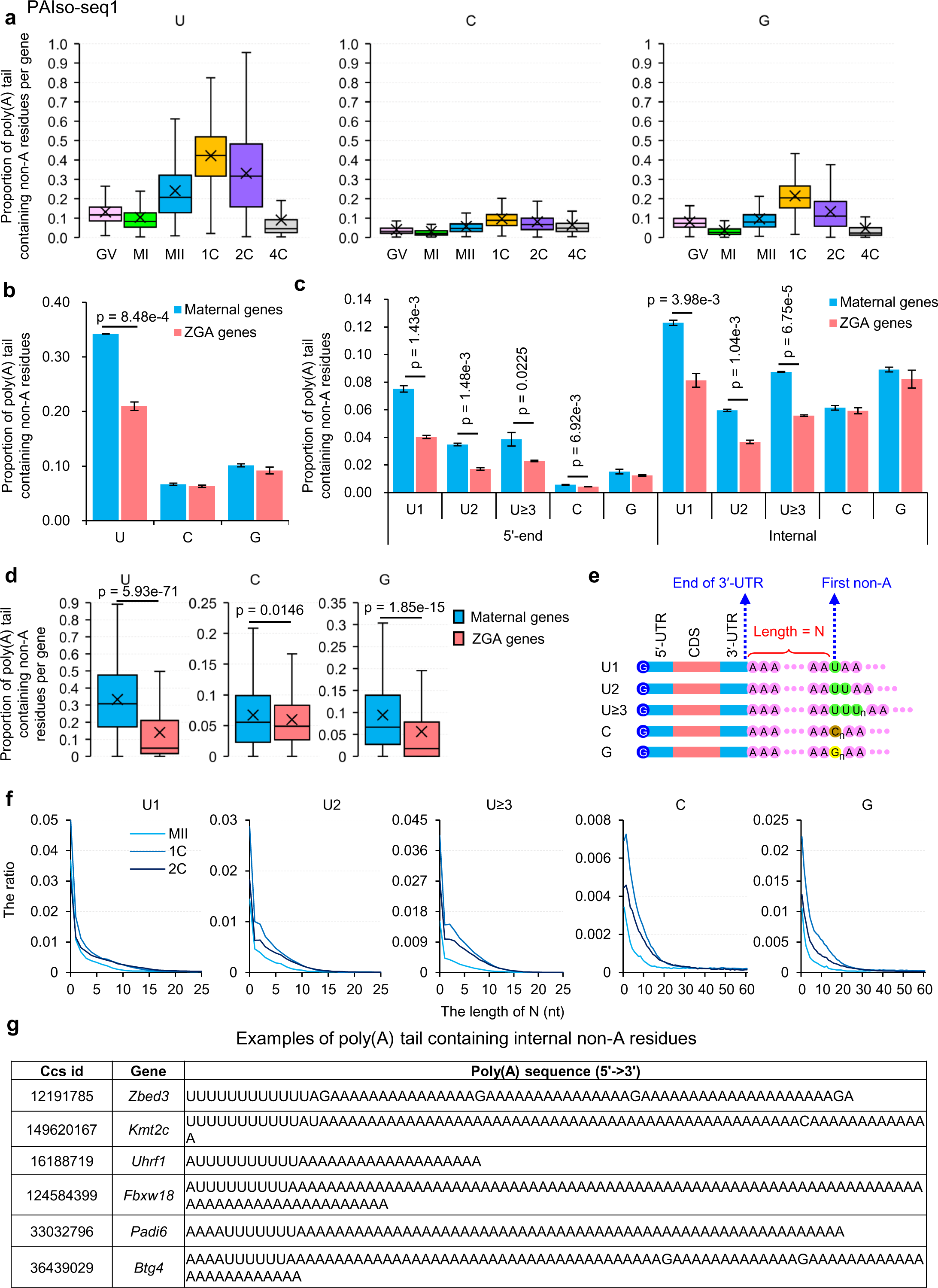
Characteristics of non-A residues during mouse OET. **a,** Box plot of the proportion of reads containing U, C, or G residues of individual genes in samples at different stages measured with PAIso-seq1. Genes (n =5,168) with at least 10 poly(A) tail containing reads (tail length ≥ 1) were included in the analysis. **b,** Overall proportion of transcripts containing U, C, or G residues in combined transcripts from maternal (n = 1,797) or zygotic genes (n = 1,448) in 2C embryos measured with PAIso-seq1. **c,** Proportion of transcripts containing 5′-end and internal U, C, or G residues for combined transcripts from maternal (n = 1,797) or zygotic genes (n = 1,448) in 2C embryos measured by PAIso-seq1. U residues were further divided according to the length of the longest consecutive number of Us (1, 2, and ≥ 3). **d,** Box plot of the proportion of reads containing U, C, or G residues for each of the maternal genes (n = 537) or zygotic genes (n = 930) in 2C embryos measured by PAIso-seq1. Genes with at least 20 poly(A) tail containing reads (tail length ≥ 1) were included in the analysis. **e,** Diagram depicting mRNA with 5’-end or internal non-A residues. N represents the length of residues between the end of 3’ UTR and the first base of the longest consecutive sequence of U, C, or G residues in a poly(A) tail. **f,** Histogram of the length of N and the ratio of U1, U2, U≥3, C, and G residues in MII, 1C, and 2C measured with PAIso-seq1. Histograms (bin size = 1 nt) are normalized to the total number of transcripts with poly(A) tails at least 1 nt in length. **g,** Examples of poly(A) tails with 5’-end or internal consecutive U residues in 1C embryos measured with PAIso-seq1. For all box plots, the “×” indicates the mean value, black horizontal bars show the median value, and the top and bottom of the box represent the values of the 25^th^ and 75^th^ percentiles, respectively. Error bars represent the SEM from two replicates. All *p* values were calculated using Student’s *t* tests.

We measured the length between the end of the 3’ UTR and the longest consecutive sequences of U, C, or G residues in poly(A) tails and recorded this as the N number (which can be as low as 0) for these poly(A) tails (Fig. 2e). The N number was short for all non-A residues (Fig. 2f, Extended Data Fig. 2f). Several poly(A) tail examples with U residues in the mouse 1C PAIso-seq1 samples all showed short N (Fig. 2g). These short N (with high levels of tails with value of 0) suggested that the non-A residues are added to short or fully deadenylated tails. Therefore, the N number is likely regulated by the deadenylation event.

### Inhibition of maternal mRNA deadenylation results in longer N in mouse MII oocytes

We reasoned that if deadenylation is impaired, longer N would be expected for poly(A) tails with non-A residues, because the substrate for adding non-A residues and re-polyadenylation is greater with longer poly(A) tails. To test this, we performed successful siRNA-mediated KD of *Btg4*, *Tut4/7*, and *Tent4a/b*, which resulted in high quality PAIso-seq1 data from these samples (Extended Data Fig. 3). *Btg4* encodes an adaptor for the CCR4–NOT deadenylase^20–22^. Tut4/7 are non-canonical poly(A) polymerases (ncPAPs) able to add U residues to short or fully deadenylated poly(A) tails^4^. Tent4a/b are ncPAPs capable of incorporating a mix of G and C residues into poly(A) tails^5^. The results showed that the N for U, C, and G residues was increased when *Btg4* was knocked-down, but not when *Tut4/7* or *Tent4a/b* was knocked-down (Fig. 3a). These observations are consistent with our findings that *Btg4* KD leads to failed deadenylation of short poly(A) tails in mouse oocytes^31^. These relatively longer tails served as substrate for re-polyadenylation, leading to longer N compared to those of short or fully deadenylated tails. Moreover, we observed a similar effect of longer N in poly(A) tails with non-A residues in human zygotes after *BTG4* KD^30^. These observations are also consistent with our findings of reduced re-polyadenylated degradation intermediates after *Btg4* KD in mice and humans^28^.

**Fig. 3.**
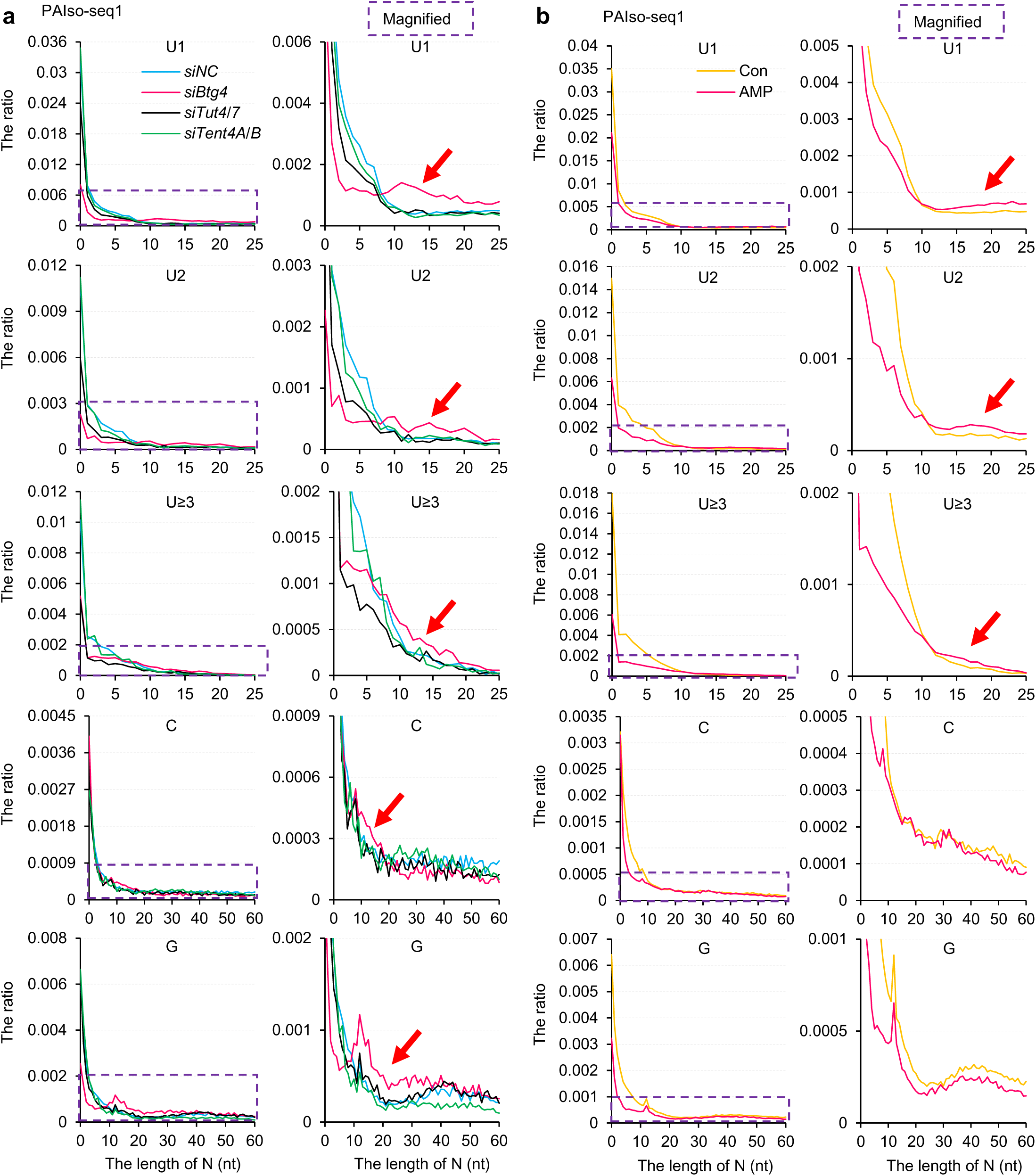
Inhibition of maternal mRNA deadenylation results in longer N in mouse MII oocytes. **a,** Histogram of the length of N and the ratio of U1, U2, U≥3, C, and G residues (from top to bottom) in *siNC*, si*Btg4*, *siTut4/7*, or *siTent4a/b* KD oocytes measured with PAIso-seq1. Histograms (bin size = 1 nt) are normalized to the total number of transcripts with poly(A) tails at least 1 nt in length. Magnified views of the regions in the purple dotted squares are shown on the right. Red arrows highlight the proportion of transcripts with increased N with *Btg4* KD. **b,** Histogram of the length of N and the ratio of U1, U2, U≥3, C, and G residues (from top to bottom) in samples with or without AMP treatment measured with PAIso-seq1. Histograms (bin size = 1 nt) are normalized to the total number of transcripts with a poly(A) tail at least 1 nt in length. Magnified views of the regions in the purple dotted squares are shown on the right. Red arrows highlight the proportion of transcripts with increased N with AMP treatment.

To further confirm this observation, we performed adenosine monophosphate (AMP) treatment of the GV oocytes and allowed them to mature *in vitro* to the MII stage (Extended Data Fig. 4). AMP can inhibit the general active deadenylation of poly(A) tails by powerfully inhibiting the major de-adenylating enzymes Cnot6l and Cnot7^32^. Consistently, we found longer N in AMP-treated MII oocytes for poly(A) tails with U residues as measured by PAIso-seq1, although the increase in N was smaller than that seen in *Btg4* KD MII oocytes (Fig. 3b). We also found that N was minimally affected for poly(A) tails with C and G residues after AMP treatment, suggesting that the effect of AMP treatment largely overlaps with *Btg4* KD, as well as an AMP-specific effect.

**Fig. 4.**
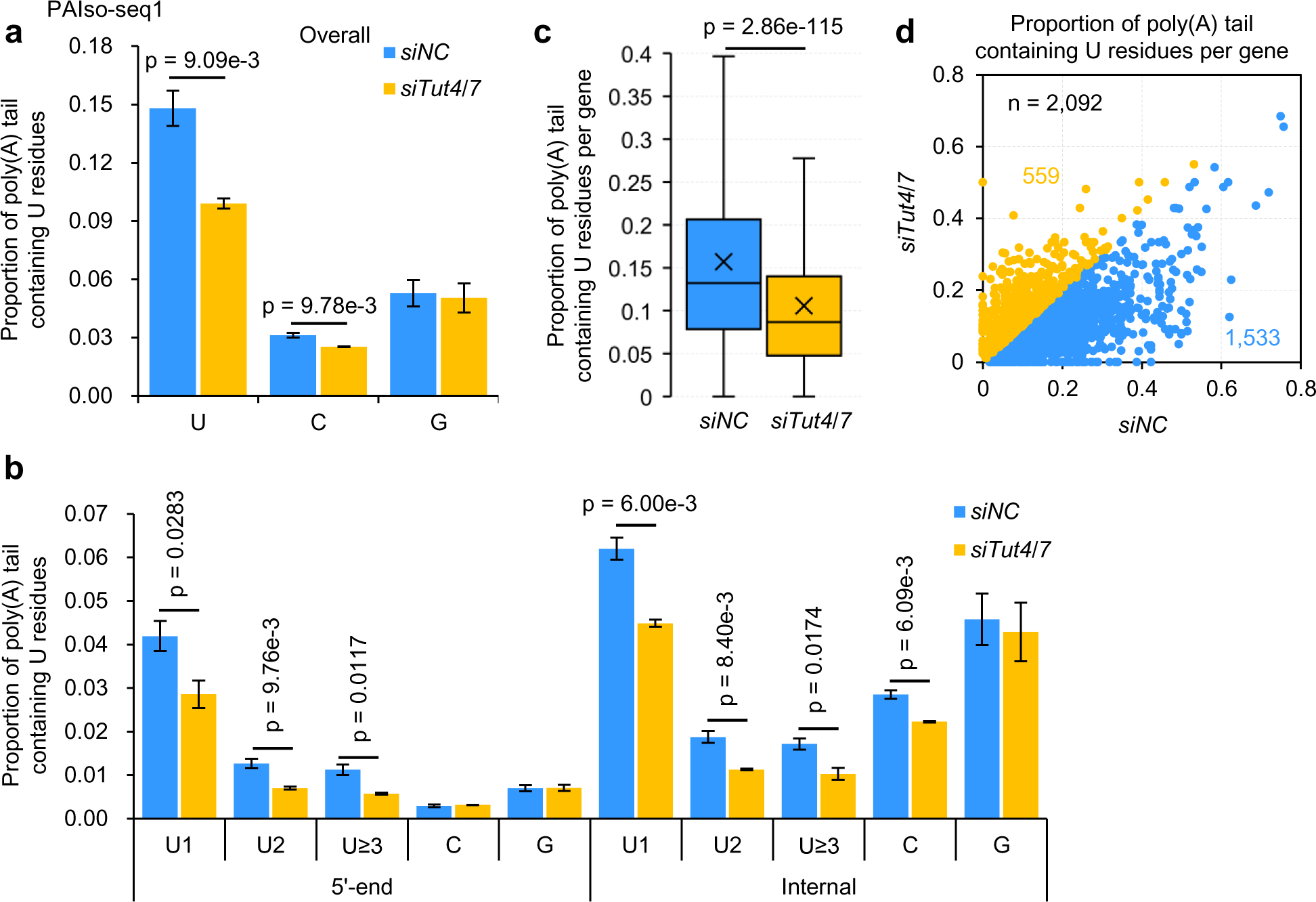
Tut4/7 is responsible for the incorporation of 5’-end and internal U residues in mouse MII oocytes. **a,** Overall proportion of transcripts containing U, C, or G residues in *siNC* and *siTut4/7* KD oocytes measured with PAIso-seq1. **b,** Proportion of transcripts containing 5′-end and internal U, C, or G residues in *siNC* and *siTut4/7* KD oocytes measured with PAIso-seq1. **c,** Box plot of the proportion of reads containing U residues of individual genes (n = 2,092) in *siNC* and *siTut4/7* KD oocytes measured with PAIso-seq1. The “×” indicates the mean value, black horizontal bars show the median value, and the top and bottom of the box represent the values of the 25^th^ and 75^th^ percentiles, respectively. **d,** Scatter plot for the proportion of reads containing U residues of individual genes in *siNC* and *siTut4/7* KD oocytes measured with PAIso-seq1. Each dot represents one gene. The poly(A) tail length for each gene is the geometric mean length of all transcripts with poly(A) tails at least 1 nt in length for the given gene. Genes with at least 20 reads in both of the samples are included in the analysis. The number of genes included in the analyses is indicated at the top left of the graph. Transcripts with a poly(A) tail of at least 1 nt were included in the analysis. Error bars represent the SEM from two replicates. All *p* values were calculated with Student’s *t* tests.

Together, these results confirmed that 5’-end and internal non-A residues are synthesized by non-A residues addition to deadenylated poly(A) tails, which are generated through Btg4-mediated global deadenylation.

### Tut4/7 is responsible for the incorporation of 5’-end and internal U residues in mouse MII oocytes

Tut4/7 can add U residues at the 3’-ends of short poly(A) tails^4, 11–13^. Our above results suggest that 5’-end and internal non-A residues are added to deadenylated tails, followed by re-polyadenylation. Therefore, we hypothesized that Tut4/7 could add U residues to deadenylated tails, which are subsequently re-polyadenylated to generate 5’-end and internal U residues. Analysis of the poly(A) tails of *Tut4/7* KD MII oocytes clearly indicated that Tut4/7 is required for the synthesis of 5’-end and internal U residues (Fig. 4a, b). At the individual gene level, the proportion of transcripts containing U residues decreased as expected in *Tut4/7* KD MII oocytes (Fig. 4c, d). A similar observation of reduced 5’-end and internal U residues has also been seen in *TUT4/7* KD human zygotes^30^. Together, these results reveal that Tut4/7 is responsible for the incorporation of 5’-end and internal U residues during the mouse OET. These 5’-end and internal U residues which are added before re-polyadenylation may have specific functions later in development.

### Tent4a/b stabilizes the re-polyadenylated mRNA by incorporating G residues in mouse MII oocytes

Tent4a/b can incorporate G residues, which can stabilize the mRNA by inhibiting the deadenylation complexes^5^. We observed incorporation of a high abundance of G residues during the mouse OET. The levels of G residues in the internal parts of poly(A) tails were much higher than those at the 5’-ends and 3’-ends during the OET; therefore, we did not separate the G residues according to the relative positions and analyzed them as a whole. Therefore, we tested whether Tent4a/4b contributed to G residues during the mouse OET. The level of G residues decreased significantly after *Tent4a/b* KD in MII oocytes (Fig. 5a). This was also evident at the individual gene level, where the proportion of transcripts containing G residues decreased after *Tent4a/b* KD (Fig. 5b, c). As G residues incorporated by Tent4a/b can stabilize the corresponding transcripts, we tested the change in transcription level upon *Tent4a/b* KD. We observed a small reduction in transcription level of all the detected genes in *Tent4a/b* KD MII oocytes (Fig. 5d). Interestingly, the transcription level of genes containing high proportion (≥30%) of transcripts with G residues in MII oocytes was greatly reduced in *Tent4a/b* KD MII oocytes (Fig. 5e), suggesting important roles of Tent4a/b in the mouse OET. Consistent with the molecular changes, our unpublished result show that fertility is reduced in *Tent4a* KO female mice. The reduction of G residues or transcription level was not seen in mouse MII oocytes depleted of *Tut4/7* or *Btg4* (Fig. 5f-i). These results reveal that poly(A) tails can be stabilized by Tent4a/b-mediated G residue incorporation after re-polyadenylation in mouse MII oocytes. Consistent with a recent study, the abundance of G residues and the transcription level also decreased in *TENT4A/B* KD human zygotes^30^. Together, our results reveal that Tent4a/b can stabilize the re-polyadenylated mRNA transcripts through the importation of G residues into poly(A) tails during the mouse OET.

**Fig. 5.**
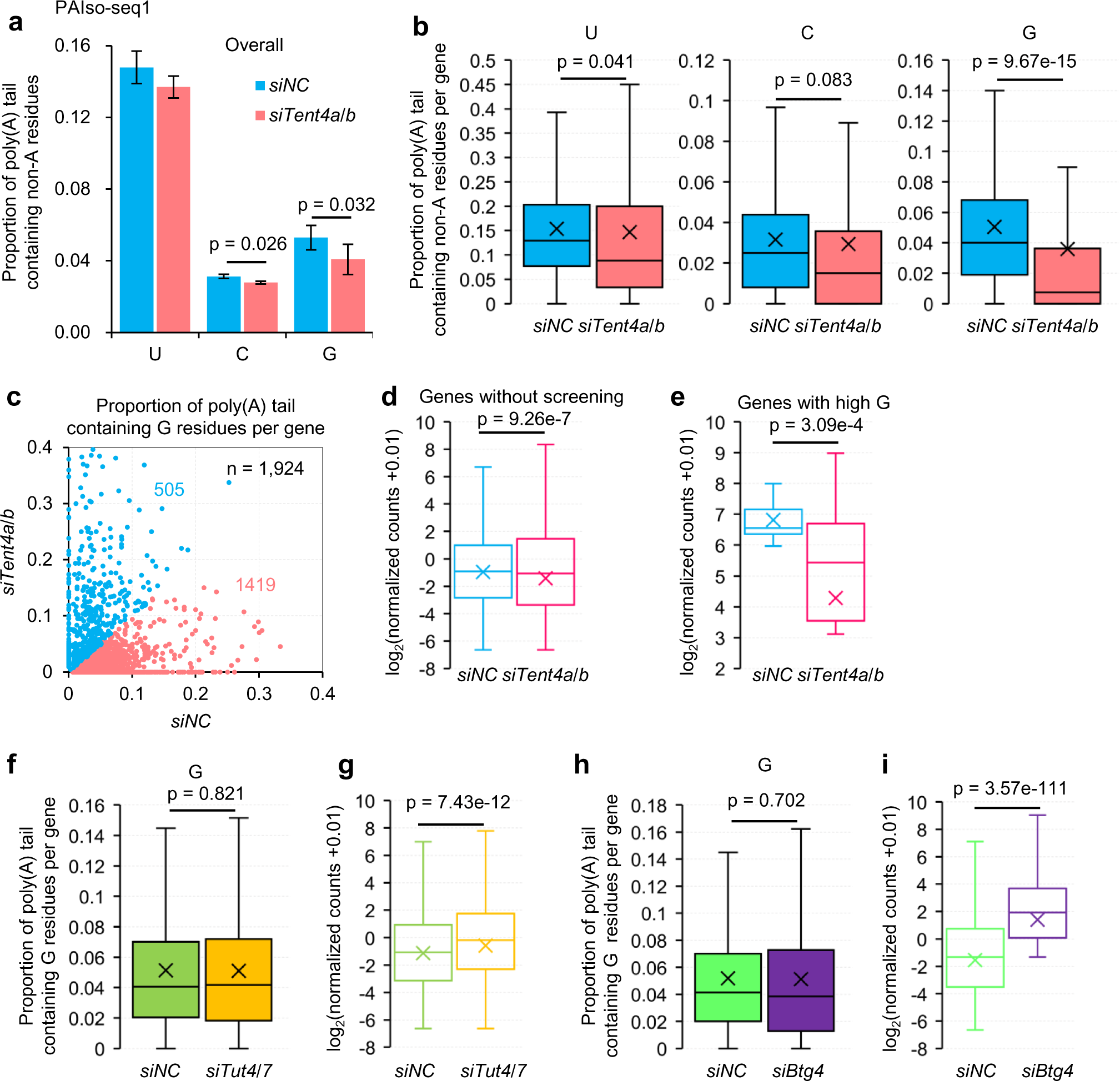
Tent4a/b stabilizes the re-polyadenylated mRNA by incorporating G residues in mouse MII oocytes. **a,** Overall proportion of transcripts containing U, C, or G residues in *siNC* and *siTent4a/b* KD oocytes measured with PAIso-seq1. **b,** Box plot of the proportion of reads containing U, C, or G residues of individual genes (n = 2,122) containing at least 20 poly(A) tails at least 1 nt in length in *siNC* and *siTent4a/b* KD oocytes measured with PAIso-seq1. **c,** Scatter plot of the proportion of reads containing G residues of individual genes in *siNC* and *siTent4a/b* KD oocytes measured with PAIso-seq1. Each dot represents one gene. The poly(A) tail length for each gene is the geometric mean length of all the transcripts with poly(A) tails at least 1 nt in length for the given gene. Genes with at least 20 reads in both of the samples were included in the analysis. The number of genes included in the analyses is indicated at the top left of the graphs. **d, e,** Box plot of the normalized counts of genes (**d**, n = 9,779) or genes with high G residues (**e**, n = 63, genes with 30% or more transcripts containing G residues) in *siNC* and *siTent4a/b* KD oocytes measured with PAIso-seq1. **f,** Box plot of the proportion of reads containing G residues of individual genes (n = 2,092) containing at least 20 poly(A) tails at least 1 nt in length in *siNC* and *siTut4/7* KD oocytes measured with PAIso-seq1. **g,** Box plot of the normalized counts of genes (n = 10,085) in *siNC* and *siTut4/7* KD oocytes measured with PAIso-seq1. **h,** Box plot of the proportion of reads containing G residues of individual genes (n = 1,957) containing at least 20 poly(A) tails of length at least 1nt in *siNC* and *siBtg4* KD oocytes measured with PAIso-seq1. **i,** Box plot of the normalized counts of genes (n = 10,907) in *siNC* and *siBtg4* KD oocytes measured with PAIso-seq1. All *p* values were calculated with Student’s *t* tests. For all box plots, the “×” indicates the mean value, horizontal bars show the median value, and the top and bottom of the box represent the values of the 25^th^ and 75^th^ percentiles, respectively. Transcripts with poly(A) tails at least 1 nt in length were included in all the analysis of proportion of reads containing non-A residues. The read counts were normalized by counts of reads mapped to protein-coding genes in the mitochondria genome if normalization was indicated.

### Maternal transcripts with U residues degrade faster after the first cleavage

Tut4/7-mediated 3’-end uridylation is important for mRNA degradation during oogenesis^12^. Maternal mRNA degradation is impaired when siRNA targeting *Tut4/7* is injected into mouse zygotes^14^. However, the function of the U residues added during oocyte maturation, which is after the GV stage and before fertilization, remains unexplored. Here, we reveal that more than 40% of mRNA transcripts contain 5’-end or internal U residues in MII oocytes and zygotes, the function of which is completely unknown. The level of poly(A) tails with U residues starts to decrease in 2C mouse embryos. Therefore, we examined the degradation speed of maternal mRNA with or without U residues in 1C and 2C mouse embryos. By separating the mRNA transcripts of each maternal gene into two categories (with or without U residues), we found that transcripts with U residues decayed faster from the 1C to the 2C stage than those without U residues, as confirmed by both the PAIso-seq1 and the PAIso-seq2 data (Fig. 6a, c). In other words, the ratio of number of transcripts with U residues to those without U residues for each maternal gene decreased from the 1C to the 2C stage (Fig. 6b, d), further confirming that the mRNA with U residues degraded faster in mouse 2C embryos.

**Fig. 6.**
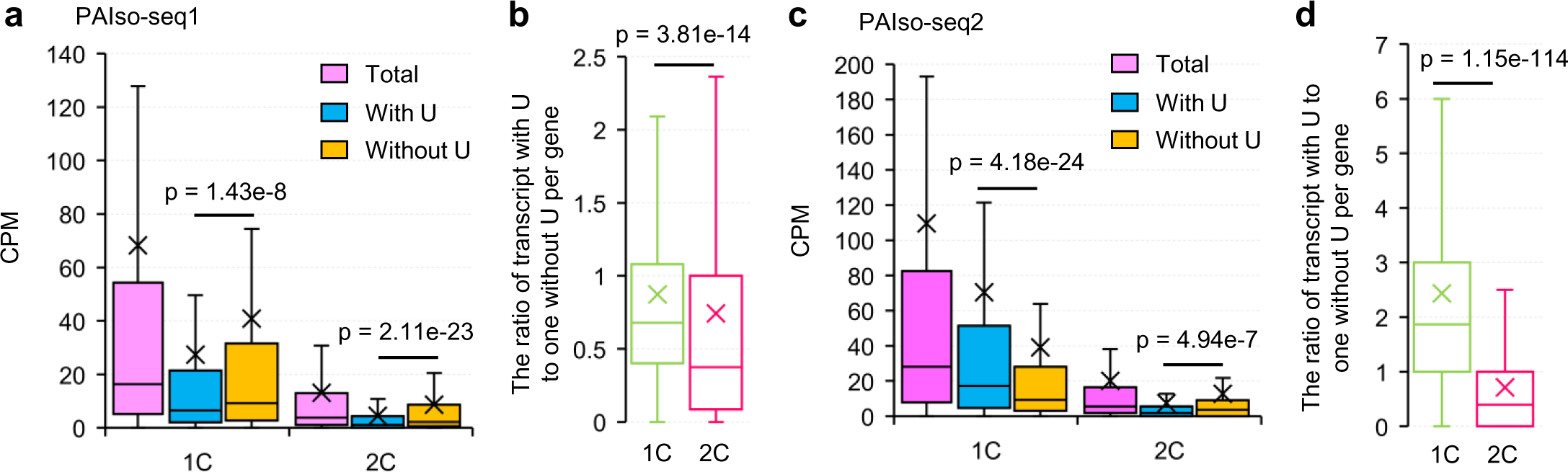
Maternal transcripts with U residues degrade faster than those without U residues. **a, c,** Box plot of the level of total transcripts, transcripts with U residues, and transcripts without U residues for each maternal gene in 1C and 2C embryos measured with PAIso-seq1 (**a,** n = 1,797) or PAIso-seq2 (**c,** n = 1,813). Transcripts with poly(A) tails at least 1 nt in length are included in the analysis. Transcript level is presented as the count per million total counts (CPM). **b, d,** Box plot of the ratio of number of transcripts with U residues to those without U residues for each maternal gene in 1C and 2C embryos measured with PAIso-seq1 (**b,** n = 1,797) or PAIso-seq2 (**d,** n = 1,813). All *p* values were calculated with Student’s *t* tests. For all box plots, the “×” indicates the mean value, horizontal bars show the median value, and the top and bottom of the box represent the values of the 25^th^ and 75^th^ percentiles, respectively.

Similarly, we also found that U residues promoted faster degradation of maternal mRNA in human 8C embryos, rat 2C embryos and pig MO embryos^29, 30^. These results reveal that mRNA with 5’-end and internal U-residues is stable when synthesized, that the 5’-end and internal U-residues mark the maternal mRNA to be degraded faster, and that degradation is activated in 2C embryos, which is an essential step in completion of the mouse OET. Tut4/7 can promote maternal mRNA degradation through addition of 3’-end U residues during oogenesis (before the OET) and after ZGA (after the OET)^12, 14^. Our finding that Tut4/7-mediated 5’-end and internal U residues serve as marks for delayed faster maternal mRNA degradation during the OET added a critical and unique puzzle piece to the role of Tut4/7 in all stages of the reproductive process.

## Discussion

The mammalian OET provides a straightforward system for studying non-A residues in poly(A) tails without complications arising from new transcription^15, 19^. In this study, we discovered that non-A residues are highly abundant and dynamic around the time of fertilization in mice (Fig. 7a). These non-A residues regulate the fate of maternal mRNA. Btg4-dependent deadenylation provides the substrate for the addition of non-A residues with U and G residues added by Tut4/7 and Tent4a/b, respectively (Fig. 7b-d). Maternal mRNA degrades quickly in 2C mouse embryos. We observed that if maternal mRNA deadenylation is blocked by treatment with AMP, maternal mRNA with U residues accumulated in the 2C AMP-treated embryos (Extended Data Fig. 5a-e). Interestingly, the ZGA as well as pre-implantation development were heavily impaired under these conditions (Extended Data Fig. 5f-h). This is consistent with the observed failed pre- implantation development of embryos derived from *Btg4* KO oocytes, due to accumulation of high-level of maternal RNA^20–22^. Therefore, correct timing of maternal mRNA clearance is essential for embryo development. About 40% of mRNA contains U residues in the 5’- end or internal parts of poly(A) tails in mouse zygotes.

**Fig. 7.**
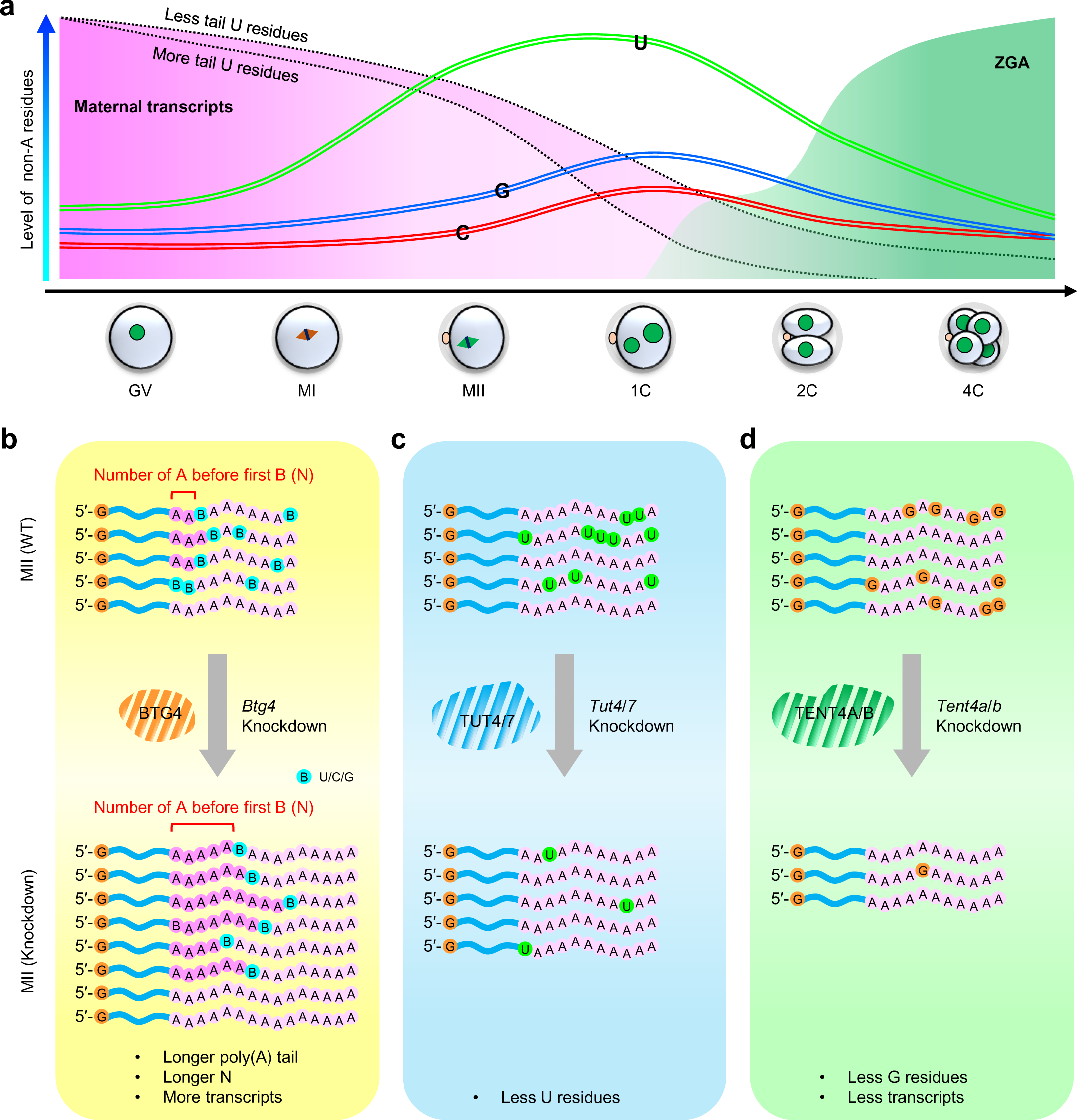
Summary of non-A residues dynamics during mouse OET. **a,** Illustration of dynamics of maternal transcripts, ZGA transcripts, and non-A (U, C, and G) residues during the mouse OET. The U residues mark maternal transcripts for rapid degradation in 2C mouse embryos. **b-d,** Illustration of the effects of *Btg4* (**b**), *Tut4/7* (**c**), or *Tent4a/b* (**d**) knockdown on the metabolism of poly(A) tails and the stability of mRNA transcripts.

Depletion of Tut4/7 during oogenesis and after fertilization both lead to failure of pre- implantation development, demonstrating the critical roles of Tut4/7-mediated uridylation for mouse reproduction^12, 14^. 3’-end U residue addition is coupled to rapid mRNA degradation in somatic cells^4, 13^. However, our study revealed that mRNA with U residues added by Tut4/7 can be temporally decoupled from immediate degradation, which can be re-polyadenylated to generate mRNA with 5’-end or internal U residues. These mRNA transcripts are stable, while the U residues in the tails mark these transcripts for rapid degradation in 2C mouse embryos for the zygotic program to take over the development process. Similarly, PIWI/piRNA does not affect the stability of its target mRNAs at the early stage of spermatogenesis, while accelerating degradation at later stages through stage-specific interaction proteins^33^. Identifying the stage-specific biochemical mechanisms responsible for the decoupling of the degradation of mRNA with U residues presents a compelling direction for future studies.

Previous studies of the dynamics and regulation of the poly(A) tail during the OET primarily come from *Drosophila*. Global poly(A) tail elongation during late oogenesis through Wispy, a Gld-2 type ncPAP, is an essential event promoting global translation in fly oocytes, while minimal changes to poly(A) tails were observed after egg activation^34–37^. Maternal mRNA typically undergoes shortening of the poly(A) tails during mouse oocyte maturation. The maternal mRNA can then be globally re-polyadenylated on short or fully deadenylated tails^28^, revealing a new type of global cytoplasmic re-polyadenylation event. Tent2/GLD2 in vertebrates can also catalyze cytoplasmic polyadenylation^38, 39^. It was therefore hypothesized that Tent2 in mice could catalyze global re-polyadenylation following fertilization, similar to the role of Wispy in flies. However, Tent2 maternal KO mice displayed normal fertility without detectable changes in the mRNA polyadenylation activity in MII-stage oocytes, which does not support Tent2 as the enzyme responsible for global re-polyadenylation after fertilization^40^. In the current study, we revealed that Tut4/7 can incorporate U residues into maternal mRNA, while Tent4a/b can incorporate G residues to stabilize the re-polyadenylated mRNA. The fertility reduction in *Tent4a* KO female mice suggests an important role for Tent4a/b during the OET. Future research directions should indicate which ncPAPs might be responsible for the global re-polyadenylation activity in mammals. While Tent2 is not a potential candidate, Tent4a/b warrants further study.

## Materials and Methods

### Animals and collection of oocytes and embryos

CD1 (ICR) mice were purchased from Beijing Vital River Laboratory Animal Technology Co., Ltd. All mouse procedures were performed in compliance with the guidelines of the Animal Care and Use Committee of the Institute of Genetics and Development Biology, Chinese Academy of Sciences. GV oocytes were isolated from ovaries after injection with 10 U of pregnant mare serum gonadotropin (PMSG) (Prospec). GV oocytes were cultured *in vitro* in M16 medium to collect MI oocytes at 8 h. To obtain MII oocytes and implantation embryos, mice were injected with 10 U of PMSG and 10 U of human chorionic gonadotropin (hCG) (Prospec) at 46- to 48-h intervals. MII oocytes were isolated from the oviduct without mating after hCG at 14-16 h. 1-cell, 2-cell, and 4-cell embryos were isolated from the oviduct with mating after hCG at 18, 40, and 60 h. Oocytes and embryos were washed in M2 medium (Sigma) and then with PBS containing 0.1% BSA (PBSA) three times and were collected into RNase-free tubes.

### Drug treatment of oocytes and embryos

The AMP (Sigma, 2 mM final concentration) was dissolved in M2 or M16 medium (Sigma). GV oocytes and 1-cell embryos were cultured in a medium containing the drug or without the drug as a control. The control and drug treated oocytes or embryos were counted or collected at the indicated times.

### Microinjection of siRNA

The set of siRNAs against *Tut4*, *Tut7*, *Btg4*, and the non-targeting control siRNA were purchased from Dharmacon. The sequence information of the siRNAs is included in Extended Data Table 1. The GV oocytes or zygotes were microinjected with 5-10 pl siRNA (10 μM) and cultured for the indicated times.

### RNA isolation

Total RNA was extracted with Direct-zol RNA MicroPrep (Zymo Research) according to the manufacturer’s instructions. Briefly, oocytes or embryo were lysed directly in 500 μl or 1 ml of TRIzol reagent (Ambion) and mixed thoroughly, and 500 μl or 1 ml of 100% ethanol was added and mixed thoroughly. The mixture was transferred into a Zymo-Spin IC Column and centrifuged to capture the RNA on the column. After washing, the RNA was eluted by adding 15-40 μl RNase-free water directly to the column matrix. The elution was repeated. Prepared RNA was stored at -80°C or used immediately.

### PAIso-seq1 and PAIso-seq2 library construction

The PAIso-seq1 libraries for mouse oocytes or embryos were constructed following the PAIso-seq1 protocol starting with purified total RNA as previously described^2^. PAIso-seq2 libraries were constructed with purified total RNA following the PAIso-seq2 protocol described in another study^10^. The barcoded oligos used in PAIso-seq1 and PAIso-seq2 are included in Extended Data Table 1. The libraries were size-selected by Pure PB beads (1x beads for cDNA more than 200 bp and 0.4x beads for cDNA more than 2 kb; the two parts of the sample were combined at equal molarity for further library construction), and made into SMRTbell Template libraries (SMRTbell Template Prep Kit). The libraries were annealed with the sequencing primer and bound to polymerase, and the polymerase-bound template was bound to Magbeads and sequenced using PacBio Sequel or Sequel II instruments at Annoroad.

### Quantitative real-time RT-PCR

The total RNA was isolated from oocytes or embryos with Direct-zol RNA MicroPrep (Zymo Research) according to the manufacturer’s instructions. SuperScript II Reverse Transcriptase (Invitrogen) was used to synthesize cDNA from RNA. Quantitative PCR was performed using TB Green Premix Ex Taq (Takara) with specific primers (Extended Data Table 1). Measurements were performed with three independent biological replicates.

### PAIso-seq1 sequencing data processing

To demultiplex and extract the transcript sequence from the CCS reads, we first matched the barcodes in the CCS reads and the reverse complements of the CCS reads allowing a maximum of two mismatches or indels (insertions and deletions). CCS reads were oriented and split into multiple transcripts if multiple barcodes were matched. Then, to obtain the precise 3’ end position of the original RNA, we aligned the matched barcode to each transcript using the semi-global function “sg_dx_trace”, which did not penalize gaps at either the beginning or the end of query/barcode in parasail package ^41^ and trimmed the barcode. Finally, the 3’-adapter and the 5’-adapter of each transcript were removed. Transcripts with length greater than 50nt were retained. Clean CCS reads were used for downstream analysis.

Clean CCS reads were aligned to the reference genome using minimap2 v.217-r941^42^ with parameters “-ax splice -uf --secondary=no -t 40 -L --MD --cs --junc-bed mm10.junction.bed”. The mm10.junction.bed file was converted from GRCm38 mouse gene annotation with “paftools.js gff2bed” in the minimap2 package. Read counts of each gene and gene assignments of each CCS reads were summarized by featureCounts v2.0.0^43^ with the parameters “-L -g gene_id -t exon -s 1 -R CORE -a Mus_musculus.GRCm38.92.gtf” using the read alignments generated by minimap2. The clean CCS reads were then ready for downstream analysis.

### PAIso-seq2 data pre-processing

The PAIso-seq2 data generated in this study was processed following previously described methods^10^, and the same initial methods used to process PAIso-seq1 data. Demultiplexing and extraction of transcript sequences followed the same procedure as for the PAIso-seq1 data, from barcode matching to trimming. For PAIso-seq2 data, we used the following regular pattern “(AAGCAGTGGTATCAACGCAG){e<=2}(AGTAC){s<=1}([ATCG]{8,12})(ATGGG) {s<=1}” to match the 5’-adapter and extract the UMIs in each transcript. Finally, the 3’-adapter and the 5’-adapter of each transcript were removed and the remaining extracted sequences were considered clean CCS, and were used for downstream analysis. Alignment of the clean CCS reads to the reference genome and gene assignment were performed the same way as in the PAIso-seq1 data pre-processing. Clean CCS reads with the identical mapping position (namely, the start and end position mapped to the reference genome) and the identical UMI (Unique Molecular Identifiers) sequence were determined, and only one clean CCS read was kept. These clean CCS reads were used for downstream analysis.

### Poly(A) tail sequence extraction

Clean CCS reads were aligned to the mouse reference genome (mm10) using minimap2 (v.217-r941) with the following parameters “-ax splice -uf --secondary=no -t 40 -L --MD -- cs --junc-bed mm10.junction.bed”^42^. Alignments with the “SA” (supplementary alignment) tag were ignored. The terminal clipped sequence of the CCS reads in the alignment bam file was used as candidate poly(A) tail sequence. We defined a continuous score based on the transitions between the two adjacent nucleotide residues throughout the 3’-soft clip sequences. To calculate the continuous score, a transition from one residue to the same residue was scored as 0, and a transition from one residue to a different residue scored as 1. The number of A, U, C, and G residues was also counted in the 3’-soft clip sequences of each alignment. The 3’-soft clip sequences with frequencies of U, C, and G all greater or equal to 0.1 were marked as “HIGH_TCG” tails. The 3’-soft clips which were not marked as “HIGH_TCG” and with continuous scores less than or equal to 12 were considered poly(A) tails.

### Poly(A) tail length measurement

To accurately determine the lengths of poly(A) tails, we only quantified the poly(A) tail length from clean CCS reads with at least ten passes. The poly(A) tail length of a transcript was calculated as the length of the sequence, including U, C, or G residues if present. The poly(A) tail length of a gene was represented by the geometric mean of the poly(A) tail length of transcripts with tail length at least 1 nt from the given gene, because poly(A) tail length distribution of a gene follows a lognormal-like distribution^35^.

### Detection of non-A residues in poly(A) tails

To minimize errors introduced by the sequencer, we used clean CCS reads with at least ten passes to identify non-A residues in poly(A) tails. G, C, and U (presented as T in CCS reads) were counted in the poly(A) tail of each CCS read. The percentage of non-A transcripts (CCS reads which contained any non-adenosine residues) of a gene was calculated as the number of CCS reads containing at least one G, C, or U residue divided by the total number of CCS reads derived from the gene. Oligo-U (U≥3) refers to reads which contain at least three consecutive Us, mono-U refers to reads which contain single U but not any two consecutive Us, and U2 refers to reads which contain UU but not any three consecutive Us. For assigning the positions of U, C, or G residues in poly(A) tails, a given poly(A) tail was first scanned for 3’-end U, C, or G residues, which if present were trimmed from the sequence, then searched for 5’-end U, C, or G residues, which were also trimmed, and finally searched for internal U, C, or G residues.

For calculating the N number for U, C, or G residues, a given poly(A) tail was first searched for the longest consecutive span of U, C, or G residues. The length of sequence before this longest consecutive stretch of U, C, or G residues was considered the N number. If a given poly(A) tail contained multiple stretches of longest consecutive U, C, or G residue, then the N number for this tail could not be determined and thus it was discarded from the N number analysis.

### Quantification and differential expression analysis of PAIso-seq data

Read counts for each gene were summarized using featureCounts. The maternal and zygotic genes were defined using the PAIso-seq1 data following a published strategy with minor modifications^44^. In brief, edgeR was used for differential expression analysis^45^. The maternal genes were defined by protein coding genes showing 4-fold enrichment in GV oocytes compared to 8C embryos at the transcript level (*p* <0.05), while the zygotic genes were defined by protein coding genes showing 4-fold enrichment in 8C embryos compared to GV oocytes at the transcript level (*p* <0.05). Pearson correlations of gene expression between replicates and different samples were calculated using the cor function, and the correlation heatmap was generated using pheatmap in R.

### Genome and gene annotation

The genome sequence used in this study is from the following links. ftp://ftp.ensembl.org/pub/release-92/fasta/mus_musculus/dna/Mus_musculus.GRCm38.dna_rm.primary_assembly.fa.gz.

The genome annotation (including the nuclear encoded mRNAs, lncRNAs and mitochondria encoded mRNAs) used in this study is from the following links. ftp://ftp.ensembl.org/pub/release-92/gtf/mus_musculus/Mus_musculus.GRCm38.92.gtf.gz

## Data Availability

The ccs data in bam format from PAIso-seq1 and PAIso-seq2 experiments will be available at Genome Sequence Archive hosted by National Genomic Data Center. Custom scripts used for data analysis will be available upon request.

## Acknowledgements

We thank Tianyao He, Jing Wu and Hongxiang Liu for their technical assistance in mouse superovulation and collection of oocytes and embryos. We thank Yiwei Zhang for his technical assistance in bioinformatic analysis. We thank Lei Li for critical reading of the manuscripts. This work was supported by the National Key Research and Development Program of China (2018YFA0107001, 2017YFA0103803, and 2018YFA0107703), the Strategic Priority Research Program of the Chinese Academy of Sciences (XDA24020203), National Natural Science Foundation of China (31970588, 32170606), Natural Science Foundation of Heilongjiang province (YQ2020C003), the China Postdoctoral Science Foundation (2020M670516, 2020T130687), and the State Key Laboratory of Molecular Developmental Biology.

## Author Contributions

Yusheng Liu, Jiaqiang Wang and Falong Lu conceived the project and designed the study. Yusheng Liu constructed the library of the PAIso-seq1 and PAIso-seq2. Yusheng Liu collected mouse oocytes and embryos for PAIso-seq1. Shuang Wu collected mouse oocytes and embryos for PAIso-seq2. Yusheng Liu performed drug treatment on oocytes and embryos. Leyun Wang, Wei Li and Qi Zhou performed siRNA mediated knock-down in oocytes. Yusheng Liu performed all other experiments. Yusheng Liu, Hu Nie, Jiaqiang Wang and Falong Lu analyzed the sequencing data. Yusheng Liu and Jiaqiang Wang organized all figures. Yusheng Liu, Jiaqiang Wang and Falong Lu supervised the project. Yusheng Liu, Jiaqiang Wang and Falong Lu wrote the manuscript with the input from the other authors.

---

**Extended Data Fig. 1.**
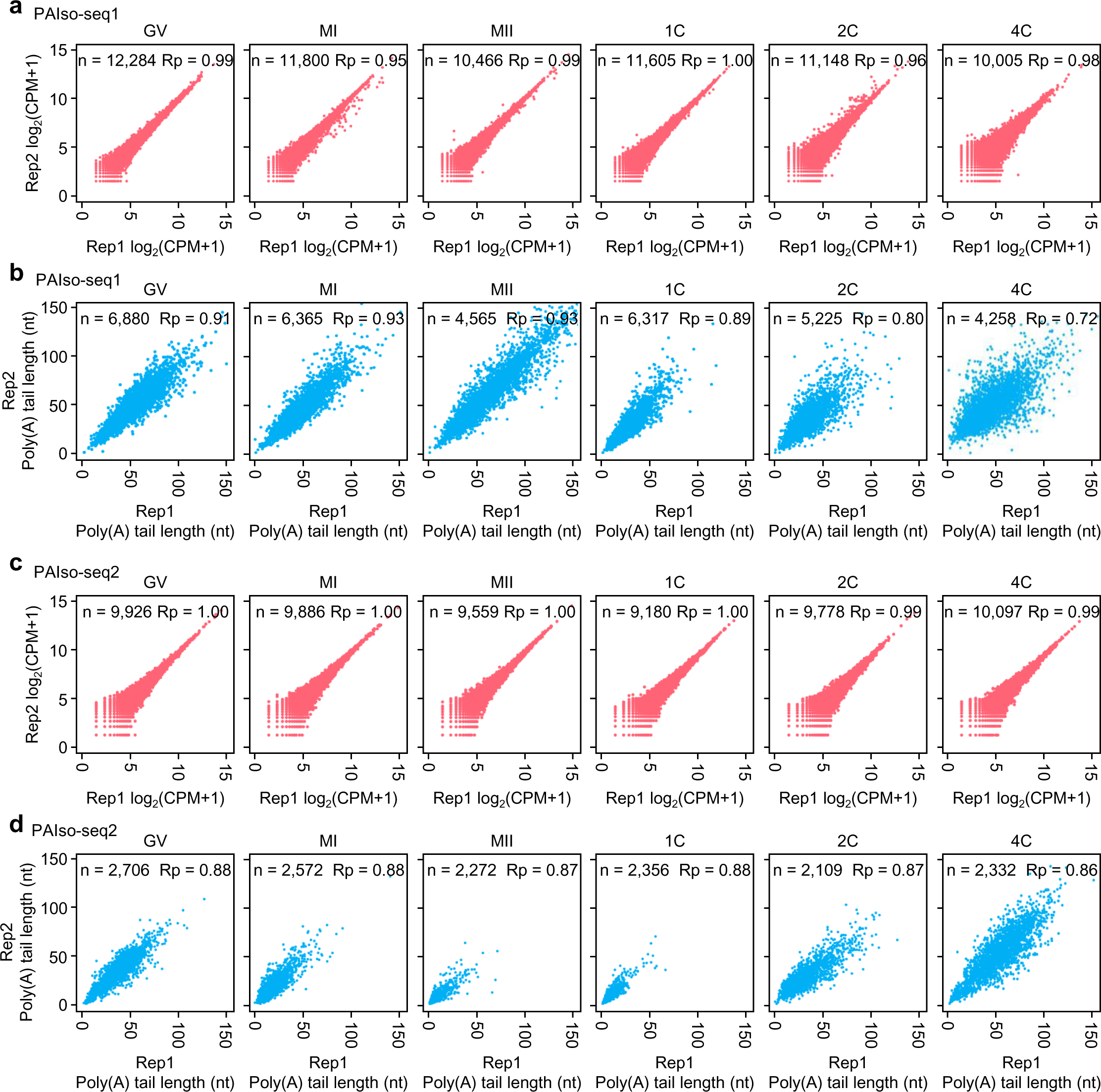
Reproducibility of PAIso-seq1 and PAIso-seq2 data for mouse oocyte and embryo samples. **a, c,** Scatter plots showing the Pearson correlation of gene expression between two replicates for mouse GV, MI, and MII oocytes, and 1C, 2C, and 4C embryos measured with PAIso-seq1 (**a**) or PAIso-seq2 (**c**). Each dot represents one gene. Pearson’s correlation coefficient (Rp) and number of genes included in the analysis are indicated at the top. **b, d,** Scatter plots showing the Pearson correlation of poly(A) tail length between two replicates for each sample measured with PAIso-seq1 (**b**) or PAIso-seq2 (**d**). Each dot represents one gene. The poly(A) tail length for each gene is the geometric mean length of all transcripts with poly(A) tails at least 1 nt in length for the given gene. Genes with at least 20 reads in each sample were included in the analysis. Pearson’s correlation coefficient (Rp) and number of genes included in the analysis are shown at the top.

**Extended Data Fig. 2.**
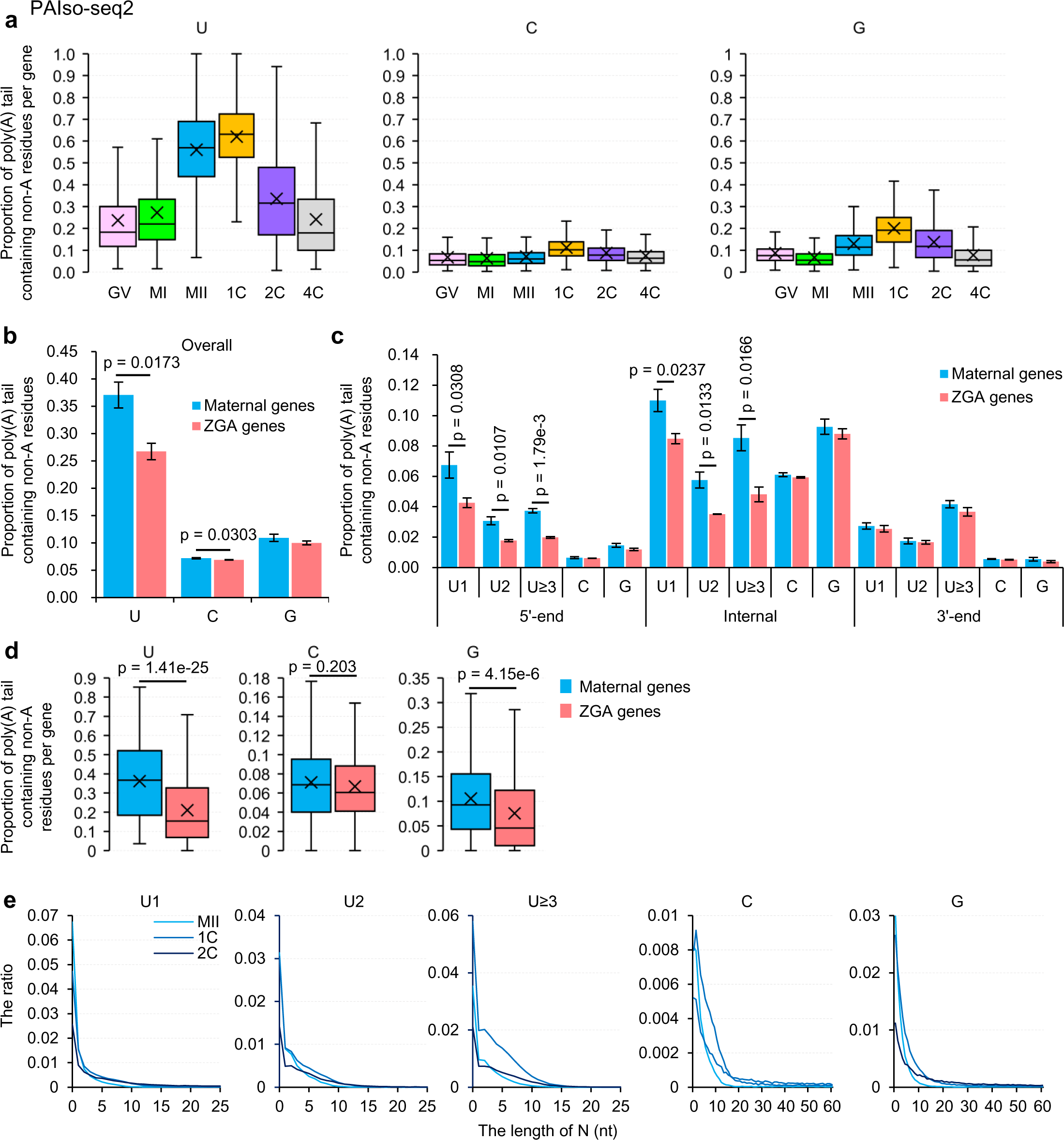
Characteristics of non-A residues incorporated during the mouse OET revealed by PAIso-seq2. **a,** Box plot of the proportion of reads containing U, C, or G residues of individual genes in samples at different stages measured with PAIso-seq2. Genes (n = 3,511) with at least 10 poly(A) tail containing reads (tail length ≥ 1) were included in the analysis. **b,** Overall proportion of transcripts containing U, C, or G residues for combined transcripts from maternal (n = 1,813) or zygotic genes (n = 1,552) in 2C embryos measured with PAIso-seq2. **c,** Proportion of transcripts containing 5′-end, internal, and 3′-end U, C, or G residues for combined transcripts from maternal (n = 1,813) or zygotic genes (n = 1,552) in 2C embryos measured with PAIso-seq2. The U residues were further divided according to the length of the longest consecutive U sequence (1, 2, and ≥3). **d,** Box plot of the proportion of reads containing U, C, or G residues for each gene of the maternal genes (n = 212) or zygotic genes (n = 740) in 2C embryos measured with PAIso-seq2. Genes with at least 20 poly(A) tail containing reads (tail length ≥ 1) were included in the analysis. **e,** Histogram of the length of N and the ratio of U1, U2, U≥3, C, and G residues in MII, 1C, and 2C measured with PAIso-seq2. Histograms (bin size = 1 nt) are normalized to the total number of transcripts with poly(A) tails at least 1 nt in length. All *p* values were calculated with Student’s *t* tests. For all the box plots, the “×” indicates the mean value, black horizontal bars show the median value, and the top and bottom of the box represent the values of the 25^th^ and 75^th^ percentiles, respectively.

**Extended Data Fig. 3.**
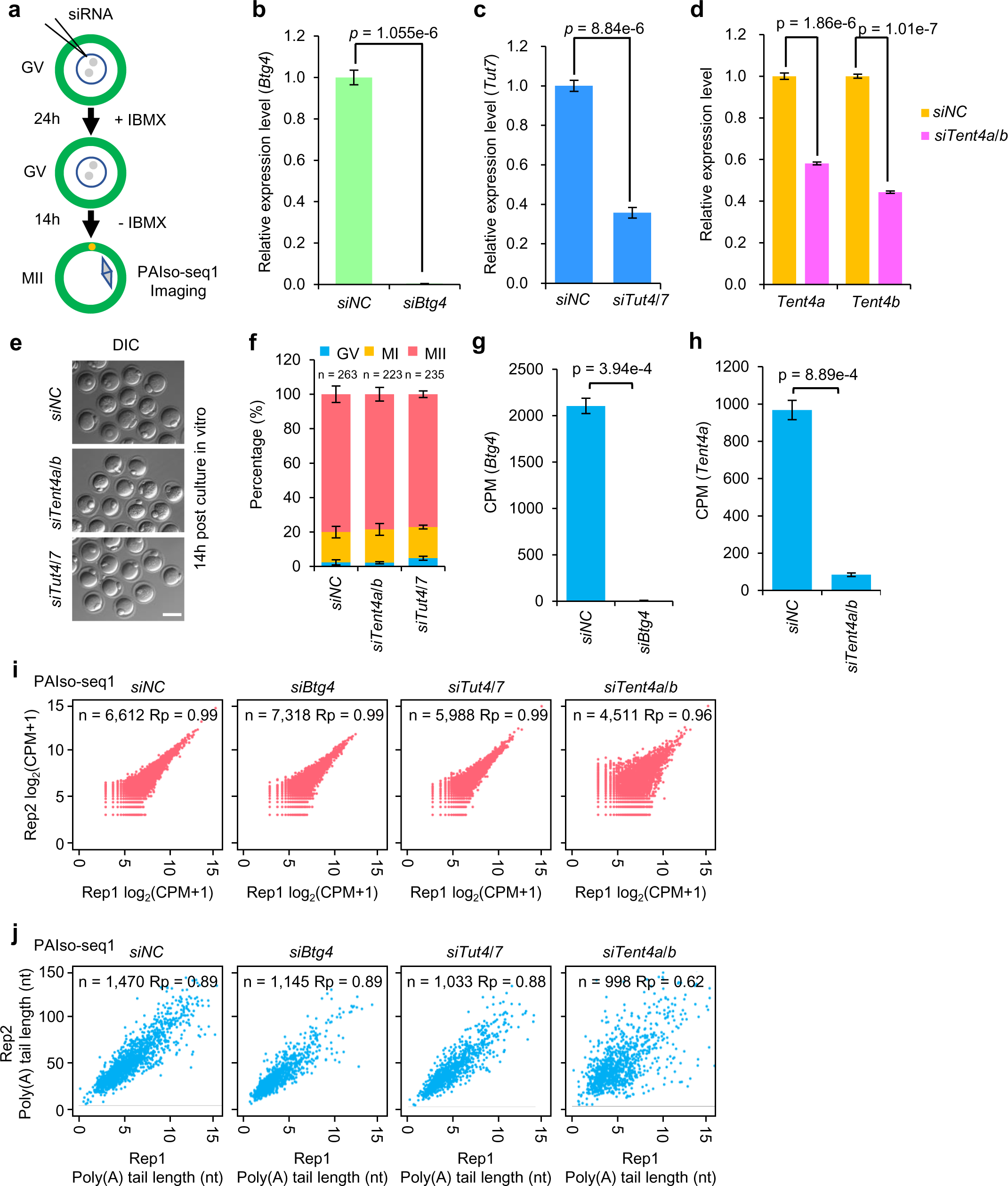
siRNA injection mediated *Btg4*, *Tut4/7*, and *Tent4a/b* knockdown in mouse oocytes. **a,** Illustration of siRNA microinjection and *in vitro* culture from the GV to MII stage. **b-d**, Verification of *Btg4* KD (**b**), *Tut7* KD (**c**), and *Tent4a/b* KD (**d**) efficiency by qPCR using samples from three biological replicates. *Tut4* expression was not detectable in control samples. Error bars indicate the SEM from two replicates. **e-f**, Morphology (**e**) and maturation rates (**f**) of control (*siNC*), *siTent4a/b*, and *siTut4/7* KD oocytes after *in vitro* maturation. Error bars indicate the SEM from two replicates. Scale bar, 100 μm. Numbers of oocytes analyzed are indicated (*siNC*, n = 263; *siTent4a/b*, n = 223; *siTut4/7*, n = 235). **g,** Expression levels of *Btg4* in *siNC* and *siBtg4* MII oocytes measured with PAIso-seq1. Error bars indicate the SEM from two replicates. **h,** Expression levels of *Tent4a* in *siNC* and *siTent4a/b* MII oocytes measured with PAIso-seq1. Error bars indicate the SEM from two replicates. **i, j,** Scatter plots showing the Pearson correlation of gene expression (**i**) and poly(A) tail length (**j**) between two replicates for *siNC*, *siBtg4*, *siTut4/7*, and *siTent4a/b* MII oocytes measured with PAIso-seq1. Each dot represents one gene. Pearson’s correlation coefficient (Rp) and number of genes included in the analysis are shown at the top. The poly(A) tail length for each gene is the geometric mean length of all transcripts with poly(A) tails at least 1 nt in length for the given gene. Genes with at least 20 reads in each sample are included in the poly(A) tail length analysis. All *p* values were calculated with Student’s *t* tests.

**Extended Data Fig. 4.**
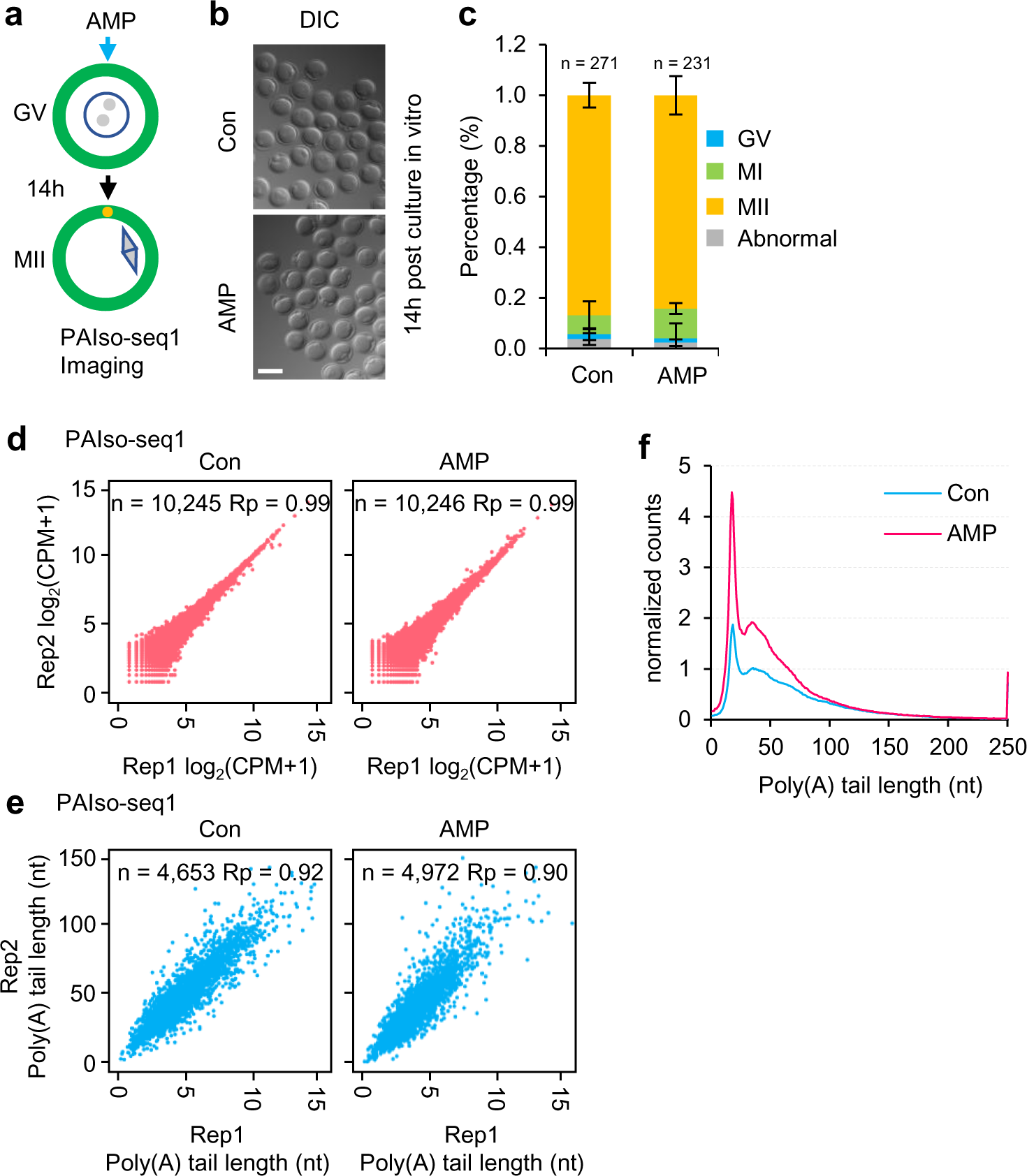
AMP treatment in mouse oocytes. **a,** Illustration of the AMP treatment experiments in mouse oocytes. **b, c**, Morphology (**b**) and maturation rates (**c**) of control (Con) and AMP-treated oocytes after *in vitro* maturation. Error bars indicate the SEM from two replicates. Scale bar, 100 μm. *p* value was calculated with Student’s *t* test. Numbers of oocytes analyzed are indicated (Con, n = 271; AMP, n = 231). **d, e,** Scatter plots showing the Pearson correlation of gene expression (**d**) and poly(A) tail length (**e**) between two replicates for control (Con) and AMP-treated MII oocytes measured with PAIso-seq1. Each dot represents one gene. Pearson’s correlation coefficient (Rp) and number of genes included in the analysis are shown at the top. The poly(A) tail length for each gene is the geometric mean length of all transcripts with poly(A) tails at least 1 nt in length for the given gene. Genes with at least 20 reads in each sample were included in the poly(A) tail length analysis. **f,** Histogram of poly(A) tail lengths of all transcripts in Control (Con), or AMP treated (AMP) mouse MII oocytes. Histograms (bin size = 1 nt) are normalized by counts of reads mapped to protein-coding genes in the mitochondria genome. Transcripts with a poly(A) tail of at least 1 nt are included in the analysis. Transcripts with poly(A) tail length greater than 250 nt are included in the 250 nt bin.

**Extended Data Fig. 5.**
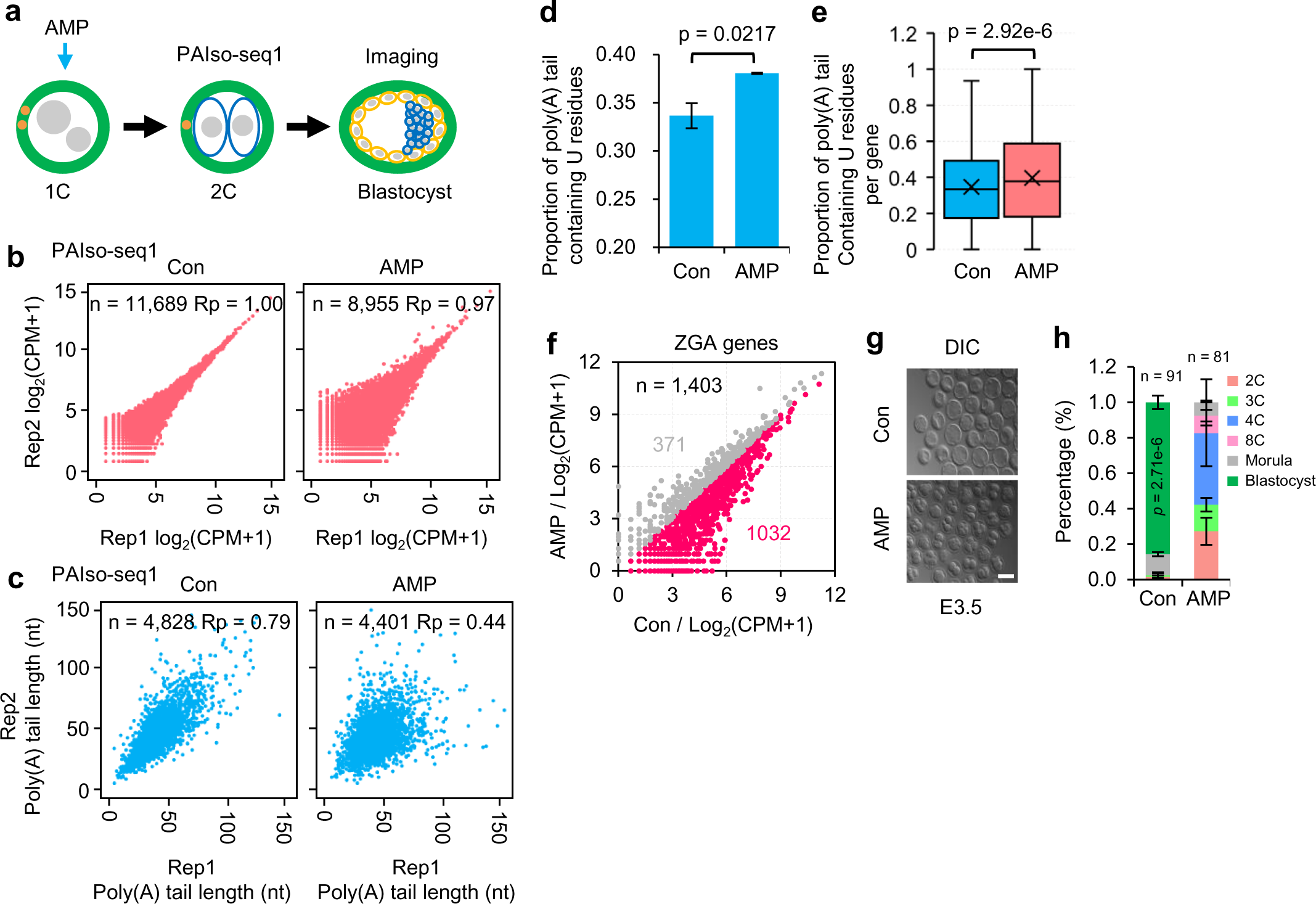
AMP treatment in mouse embryos. **a,** Illustration of the AMP treatment experiments. **b, c,** Scatter plots showing the Pearson correlation of gene expression (**b**) and poly(A) tail length (**c**) between two replicates for control (Con) and AMP-treated 2C embryos measured with PAIso-seq1. Each dot represents one gene. Pearson’s correlation coefficient (Rp) and number of genes included in the analysis are shown at the top. The poly(A) tail length for each gene is the geometric mean length of all transcripts with poly(A) tails at least 1 nt in length for the given gene. Genes with at least 20 reads in each sample were included in the poly(A) tail length analysis. **d,** Proportion of combined transcripts from maternal genes (n = 1,797) containing U residues from mouse 2C embryos with or without AMP treatment measured with PAIso-seq1. Transcripts with poly(A) tails at least 1 nt in length were included in the analysis. **e,** Box plot of the proportion of reads containing U residues for individual maternal genes (n = 484) with at least 20 poly(A) tails at least 1 nt in length in mouse 2C embryo samples with or without AMP treatment measured with PAIso-seq1. **f,** Scatter plot of the expression level of ZGA genes in mouse 2C embryo samples with and without AMP treatment measured with PAIso-seq1. Each dot represents one gene. Genes with at least 1 read in one of the samples were included in the analysis. The number of genes included in the analyses is included at the top left of the graph. **g,** Morphology of control and AMP-treated mouse embryos at embryonic day 3.5 (E3.5). Scale bar, 100 μm. **h,** Developmental rates of control (Con) and AMP-treated mouse embryos examined at E3.5. Error bars indicate the SEM from two replicates. The numbers of analyzed embryos are indicated (n = 91 for Con, n = 81 for AMP). All *p* values were calculated with Student’s *t* tests. For all box plots, the “×” indicates the mean value, horizontal bars show the median value, and the top and bottom of the box represent the values of the 25^th^ and 75^th^ percentiles, respectively.

